# Unique genetic features of the naked mole-rat’s *THADA* gene

**DOI:** 10.1101/2021.09.19.460947

**Authors:** Khadijah Banjar, Carsten Holzmann, Jörn Bullerdiek

## Abstract

*Thyroid Adenoma Associated* (*THADA*) is a protein-coding gene that maps to chromosomal band 2p21 and first has been described as a target of recurrent translocation partner in thyroid tumors. Many genome-wide association studies have revealed an association between *THADA* and two frequent human diseases, i.e. type 2 diabetes and polycystic ovary syndrome. Nevertheless, the function of its protein is not been completely understood. However, recent evidence suggests that in a Drosophila model THADA can act as a sarco/endoplasmic reticulum Ca^2+^-ATPase (SERCA)-interacting protein which uncouples SERCA from this function. Once being uncoupled, SERCA produces an increased amount of heat without transporting calcium thus triggering nonshivering thermogenesis. This data prompted us to compare human THADA with that of 65 other eutherian mammals. This includes a comparison of THADA of a variety of eutherian mammals with that of the naked-mole rat (*Heterocephalus glaber*) which is known to display unique features of thermoregulation compared to other mammals. Our analysis revealed five positions where only the naked-mole rat presented differences. These latter positions included four single amino acid substitutions and one unique deletion of six or seven amino acids, respectively, between residues 858 and 859. In future studies these changes will be analyzed further in detail for their functional relevance.

## Introduction

Human *Thyroid Adenoma Associated* (*THADA*) maps to chromosomal band 2p21. First it has been described as a recurrent translocation partner in thyroid tumors characterized by corresponding reciprocal chromosomal translocations with a predominance of a t(2;7)(p21;p15) (Rippe et al., 2003). Later, *THADA* was found to be expressed at a high level in follicular cells of normal thyroid tissue when compared with other tissues (Kloth et al., 2011) but it was shown that its expression was not altered in thyroid cancers positive or negative for *THADA* fusions. It has been proposed that in these latter rearrangements, *THADA* drives tumorigenesis by juxtaposing its regulatory sequences to a translocation partner thereby activating a gene of this partner. In the majority of cases, the gene encoding insulin-like growth factor 2 mRNA-binding protein 3 (*IGF2BP3*) serves as this activated partner (Panebianco et al., 2017). Accordingly, translocations of *THADA* have been placed among the 20 most-significant significantly recurrent breakpoints (SRBs) observed in non-coding regions in human cancers (Rheinbay et al., 2020). Nevertheless, as a result of the translocations not only *IGF2BP3* becomes upregulated but also a truncated *THADA* is expressed. Thus, it remains to be investigated, if both effects - upregulation of *IGF2BP3* as well as the presence of truncated THADA-contribute its potential of driving tumorigenesis. As to the tumor types affected, said *THADA* rearrangements in the majority of cases characterize a particular type of thyroid neoplasms called non-invasive follicular thyroid neoplasm with papillary-like nuclear features (NIFTP) (Pool et al., 2019) but other histologic types have been reported as well. In a study by Morariu et al. (2021) thirty nodules positive for *THADA*-*IGF2BP3* fusion on FNA were detected. Of the positive nodules that were managed surgically all were intrathyroidal and all lacked aggressive histology. Surgical pathology revealed seven (32%) malignant nodules (six encapsulated follicular variant papillary thyroid carcinomas (EFVPTC), one minimally infiltrative FVPTC), ten (45%) noninvasive follicular thyroid neoplasms with papillary-like nuclear features (NIFTP), and five (23%) nodules that were classified as follicular adenomas (FA).

Despite its important role in the genesis of thyroid neoplams THADA’s normal physiological functions still remained enigmatous. Besides the thyroid *THADA* is abundantly expressed in some other endocrine tissues as pituitary gland and parathyroid gland as well as in the endometrium (https://www.proteinatlas.org/ENSG00000115970-THADA/tissue).

Many genome-wide association studies (GWAS) have revealed an association between *THADA* and two frequent human diseases, i.e. type 2 diabetes (T2DM) (e.g. Staiger et al., 2008, Grarup et al., 2008, Kang et al., 2009, Boesgaard et al., 2009, Hu et al., 2009, Simonis-Bik et al., 2010, Schleinitz et al., 2010, Cheng et al., 2011, Gupta et al., 2012, Hotta et al., 2012, Villegas et al., 2014, DeMenna et al., 2014, Bai et al., 2015, Prasad et al., 2016) and polycystic ovary syndrome (PCOS) (e.g. Zhao et al., 2012, Eriksen et al., 2012, Wang et al., 2012, Cui et al., 2013, Sun et al., 2014, Day et al., 2015, Cui et al., 2015, Xia et al., 2019, Park et al., 2019, Bakhashab et al., 2019, Tian et al., 2020, Vishnubotla et al., 2020). Again, due to a lack of knowledge, for a long time it was impossible to tie these associations with any type of causal relationship. Nevertheless, both associations turned out to be highly reproducible and apparently did not depend on ethnicity (Goodarzi et al., 2012).

A genome-wide comparison of the Neandertal genome with recent humans has identified SNPs in *THADA* as the most strongly positively selected SNPs during evolution of modern humans (Green et al., 2010). In 2014, *THADA* was marked as a cold adaptation gene based on population genetic studies (Cardona et al., 2014) but the remarkable and nearly total lack of knowledge about the function of THADA remained until 2017 when, using a *Drosophila* model, Moraru et al. (2017) were able to show that THADA can act as a sarco/endoplasmic reticulum Ca^2+^-ATPase (SERCA)-interacting protein. SERCA is a calcium transporter from the cytosol into the sarcoplasmic reticulum and its action contributes to obligative energy expenditure. Apparently, like sarcolipin, THADA is able to uncouple SERCA from this function and, once being uncoupled, SERCA produces an increased amount of heat without transporting calcium thus triggering nonshivering thermogenesis (Chatterjee et al., 2017). Nevertheless, unlike sarcolipin THADA does not seem to be inducible by cold, pointing to a more complex network. The *Drosophila* experiments suggest that *THADA* is not only a very „old“ gene in terms of evolution but also has served functions in thermogenesis very early. These functions may explain the association between *THADA* and T2DM as well as with PCOS and suggest a causal link. The anti-obesity effect of THADA may be mediated through enhanced thermogenesis or reduced ER (endoplasmic reticulum) calcium. Of note, in mice changes of calcium levels in the endoplasmic reticulum during pancreatic maturation were correlated with the expression of *THADA* (West et al., 2021). Overall, an in depth characterization of the metabolic effects of the THADA-driven network will not only aim at a better understanding of its function in general but might also provide new therapeutic targets for the treatment of both conditions associated with *THADA* by GWAS (Chatterjee et al., 2017).

*THADA* as well as its protein were found to be highly conserved among mammals and even among vertebrates (Drieschner et al., 2007, Soller et al., 2008). To better understand functional relationships between THADA and thermogenesis, this study aims at the characterization of Thada of the naked-mole rat (NMR Thada) (*Heterocephalus glaber*) which, when compared to other rodents and mammals in general, is expressing many unusual traits including its limited ability to maintain a stable body temperature and, at least in part together with other bathyergids, appears to have a different thyroid hormone physiology. While some evidence has been presented that NMR are even non-ageing mammals i.e. lack an increase of likelihood of mortality with chronological age (Ruby et al., 2018) this has been a matter of controversial debate (Dammann et al., 2019, Ruby et al., 2019). However, there is still ample data from the current literature allowing to conclude that NMR are at least characterized by a remarkably late onset of senescence (Braude et al., 2021).

## Materials and methods

THADA is a well conserved protein among mammalian species. For protein alignment of multiple sequences the NCBI blast tool has been used (https://blast.ncbi.nlm.nih.gov/Blast.cgi?PAGE=Proteins).

## Results

In a first approach, THADA AS-sequences of a small set of eutherian (placental) mammals have been compared with that of the naked-mole rat by multiple sequence alignment. For this first analysis we referred to NMR Thada sequence XP_004839417.1 (NCBI reference sequence) and compared it to a first set of 16 species including:

*Bos taurus* (European cattle), *Callithrix jacchus* (common marmoset), *Canis lupus familiaris* (dog), *Dasypus novemcinctus, Fukomys damarensis* (Damaraland mole-rat), *Gorilla gorilla gorilla* (gorilla), *Homo sapiens, Loxodonta africana* (African bush elephant), *Macaca mulatta* (rhesus macaque), *Mus musculus* (house mouse), *Myotis brandtii* (Brandt’s bat), *Oryctolagus cuniculus* (European rabbit), *Otolemur garnettii* (northern greater galago), *Pan troglodytes* (chimpanzee), *Papio anubis* (olive baboon), and *Rattus norvegicus* (Norwegian rat).

THADA was unambiguously identified in all these species. Furthermore, this search confirmed the high level of identity between human THADA and that of the other mammalian species investigated ranging between 99.1 % (gorilla and chimpanzee) and 78.3 % (mouse) (Tab.1). Between human and NMR-Thada, an identity of 85.1% was observed.

**Tab. 1:**

| ☒ select all 17 sequences selected |  | GenPept | Graphics | Distance tree of results | Multiple alignment | New MSA Viewer |  |  |  |
| --- | --- | --- | --- | --- | --- | --- | --- | --- | --- |
|  | Description | Scientific Name | Max Score | Total Score | Query Cover | E value | Per. Ident | Acc. Len | Accession |
| ☒ | thyroid adenoma-associated protein isoform a [Homo sapiens] | Homo sapiens | 4026 | 4026 | 100% | 0.0 | 100.00% | 1953 | NP_001077422.1 |
| ☒ | thyroid adenoma-associated protein [Gorilla gorilla gorilla] | Gorilla gorilla gorilla | 3992 | 3992 | 100% | 0.0 | 99.13% | 1953 | XP_004029196.2 |
| ☒ | thyroid adenoma-associated protein isoform X1 [Pan troglodytes] | Pan troglodytes | 3992 | 3992 | 100% | 0.0 | 99.13% | 1953 | XP_001141732.2 |
| ☒ | thyroid adenoma-associated protein isoform X6 [Macaca mulatta] | Macaca mulatta | 3860 | 3860 | 99% | 0.0 | 96.24% | 1942 | XP_002799292.2 |
| ☒ | thyroid adenoma-associated protein isoform X1 [Papio anubis] | Papio anubis | 3747 | 3747 | 99% | 0.0 | 96.23% | 1934 | XP_017802961.1 |
| ☒ | thyroid adenoma-associated protein isoform X1 [Callithrix jacchus] | Callithrix jacchus | 3672 | 3672 | 100% | 0.0 | 92.83% | 1952 | XP_035128574.1 |
| ☒ | PREDICTED: thyroid adenoma-associated protein isoform X1 [Myotis brandtii] | Myotis brandtii | 3336 | 3336 | 100% | 0.0 | 84.23% | 1950 | XP_014393943.1 |
| ☒ | thyroid adenoma-associated protein isoform X3 [Heterocephalus glaber] | Heterocephalus glaber | 3303 | 3303 | 98% | 0.0 | 85.09% | 1928 | XP_004839417.1 |
| ☒ | thyroid adenoma-associated protein homolog [Canis lupus familiaris] | Canis lupus familiaris | 3301 | 3301 | 99% | 0.0 | 83.35% | 1948 | NP_001103429.1 |
| ☒ | thyroid adenoma-associated protein isoform X1 [Fukomys damarensis] | Fukomys damarensis | 3299 | 3299 | 98% | 0.0 | 84.21% | 1934 | XP_033619720.1 |
| ☒ | thyroid adenoma-associated protein isoform X2 [Dasypus novemcinctus] | Dasypus novemcinctus | 3291 | 3291 | 100% | 0.0 | 84.54% | 1945 | XP_012379587.1 |
| ☒ | PREDICTED: thyroid adenoma-associated protein [Oryctolagus cuniculus] | Oryctolagus cuniculus | 3276 | 3276 | 97% | 0.0 | 85.14% | 1942 | XP_008252690.2 |
| ☒ | thyroid adenoma-associated protein isoform X1 [Bos taurus] | Bos taurus | 3253 | 3253 | 100% | 0.0 | 83.80% | 1946 | XP_010808315.1 |
| ☒ | thyroid adenoma-associated protein [Rattus norvegicus] | Rattus norvegicus | 3111 | 3111 | 100% | 0.0 | 79.13% | 1937 | NP_001178698.1 |
| ☒ | thyroid adenoma-associated protein homolog [Mus musculus] | Mus musculus | 3068 | 3068 | 100% | 0.0 | 78.36% | 1938 | NP_898842.2 |
| ☒ | thyroid adenoma-associated protein isoform X1 [Otolemur garnettii] | Otolemur garnettii | 3053 | 3053 | 90% | 0.0 | 87.12% | 1753 | XP_023367895.1 |
| ☒ | thyroid adenoma-associated protein isoform X1 [Loxodonta africana] | Loxodonta africana | 2847 | 2847 | 86% | 0.0 | 84.47% | 1683 | XP_023411086.1 |

A total of six amino acid residues of NMR-Thada turned out to be unique among orthologs present in these 16 other mammalian species with no variation of each of the corresponding amino acid seen in the other mammals analyzed. In addition, NMR-Thada differed from all other investigated mammalian species by a unique deletion site of six or seven amino acids, respectively, between residues 858 and 859 (Fig. 2).

**Legend Fig. 1:**
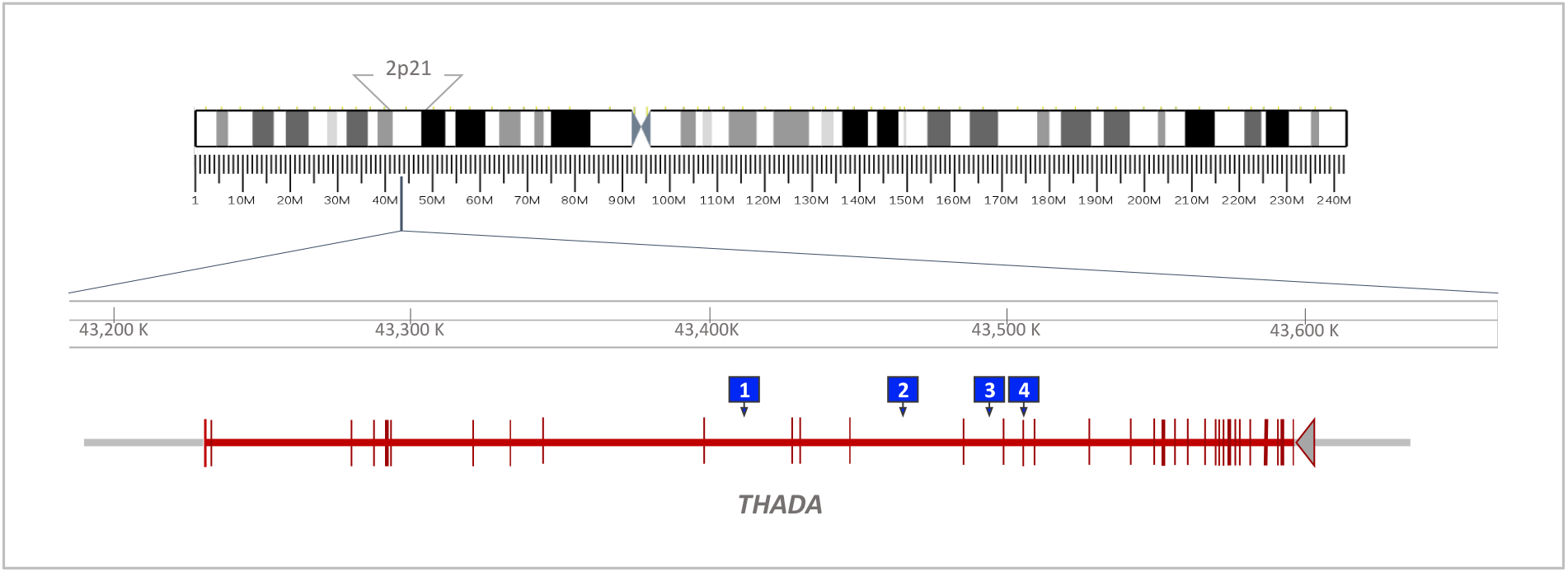
Chromosomal assignment of human *THADA* to band 2p21, intron-exon structure of *THADA* (red, exons represented by vertical bars), and localization of SNPs (blue squares) associated with Type 2 Diabetes (T2DM) and Polycystic Ovary Syndrome (PCOS), respectively. 1: rs13429458, PCOS, 2: rs10203174, T2DM, 3: rs12478601, PCOS, 4: rs7578597, T2DM

**Legend Fig. 2:**
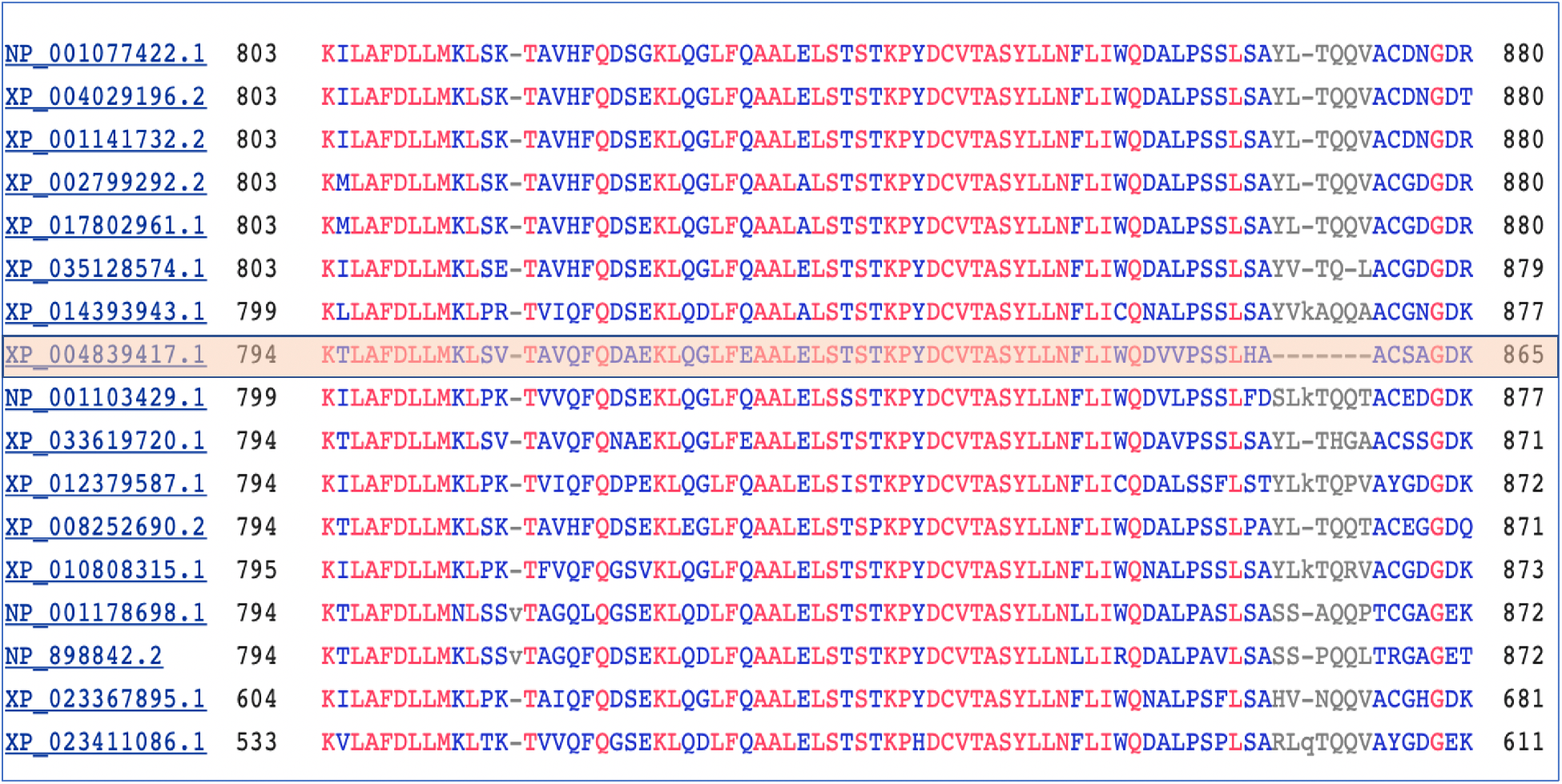
Part of human THADA (first line) aligned that of a variety of eutherian mammals including *H. glaber* (light orange) showing a unique deletion present in Thada of H. glaber. Accession numbers refer to the following species (see Tab. 1): *Homo sapiens:* NP_001077422.1, *Gorilla gorilla gorilla:* XP_004029196.2, *Pan troglodytes:* XP_001141732.2, *Macaca mulatta:* XP_002799292.2, *Papio anubis:* XP_017802961.1, *Callithrix jacchus:* XP_035128574.1, *Myotis brandtii:* XP_014393943.1, *Canis lupus familiaris:* NP_001103429.1, *Fukomys damarensis*: XP_033619720.1, *Dasypus novemcinctus:* XP_012379587.1, *Oryctolagus cuniculus:* XP_008252690.2, *Bos taurus:* XP_010808315.1, *Rattus norvegicus:* NP_001178698.1, *Mus musculus:* NP_898842.2, *Otolemur garnettii:* XP_023367895.1, *Loxodonta africana:* XP_023411086.1

As a next step, THADA of an extended second set of further 49 non-primate eutherian mammals was compared by multiple alignment with that of *H. glaber*. This second set comprised the following species:

> *Acinonyx jubatus, Balaenoptera musculus, Camelus dromedarius, Camelus ferus, Canis lupus dingo, Ceratotherium simum simum, Choloepus didactylus, Chrysochloris asiatica, Delphinapterus leucas, Eptesicus fuscus, Equus asinus, Equus caballus, Eumetopias jubatus, Felis catus, Globicephala melas, Halichoerus grypus, Ictidomys tridecemlineatus, Lagenorhynchus obliquidens, Lynx canadensis, Lynx pardinus, Manis javanica, Manis pentadactyla, Marmota flaviventris, Marmota monax, Miniopterus natalensis, Mirounga leonine, Molossus molossus, Monodon Monoceros, Myotis lucifugus, Myotis myotis, Neomonachus schauinslandi, Neophocaena asiaeorientalis, Nyctereutes procyonoides, Odobenus rosmarus divergens, Orcinus orca, Panthera pardus, Panthera tigris altaica, Phocoena sinus, Pteropus vampyrus, Puma yagouaroundi, Rhinolophus ferrumequinum, Rousettus aegyptiacus, Talpa occidentalis, Trichechus manatus latirostris, Tursiops truncates, Ursus arctos horribilis, Ursus maritimus, Vulpes Vulpes, Zalophus californianus*

Four of the previously identified six amino acid residues of NMR-Thada remained to be unique even among the extended set of mammalian species (Tab. 2). Also, the deletion as described above remained unique and represents the most obvious difference between *H. glaber* sequence XP_004839417.1 and all other mammals included in the comparison. Next, all other NMR THADA sequences as available through NCBI were also checked for these differences. In total seven Thada sequences from the naked-mole rat are currently available by NCBI (reference sequence protein isoforms for the Thada gene) and were analyzed (Tab. 3).

**Tab. 2:** Aa position refers to *H. glaber* sequence: XP_004839417.1

| **NMR Thada,**<br>**Aa position** | **NMR** | **all other mammals analyzed** |
| --- | --- | --- |
| 696 | histidine | glutamine |
| 1118 | arginine | lysine |
| 1314 | lysine | arginine |
| 1358 | leucine | phenylalanine |

**Tab. 3:** *H. glaber* Thada sequences as available through (https://blast.ncbi.nlm.nih.gov/Blast.cgi?PAGE=Proteins). *: several differences to the other clones

| sequence ID |
| --- |
| XP_004839417.1 |
| EHA97799.1* |
| XP_021109460.1 |
| XP_021109461.1 |
| XP_021109459.1 |
| XP_021109457.1 |
| XP_021109458.1 |

Since apparently important insights into the function of THADA have been obtained by in depth analysis of its counterpart in *Drosophila melanogaster* (Moraru et al., 2017), NMR-Thada was also compared with that of the fruit fly. Analysis revealed a 29.8 % identity between THADA of both species which is in the same range as that between the fly and *Homo sapiens* (29.1 % identity).

## Discussion and further experimental approach

Human THADA and its gene have been identified nearly twenty years ago by positional cloning of a breakpoint region of chromosome 2 recurrently involved in chromosomal alterations in thyroid tumors (Bol et al., 2001, Rippe et al., 2003). In genome-wide association studies, the SNPs of the *THADA* gene have been found to be associated with T2DM as well as with PCOS and SNPs in *THADA* belong to the most strongly positively selected SNPs during the evolution of the recent human species (Green et al., 2010). Nevertheless, surprisingly little is known about THADA function. Some recent data, however, suggest that, as a SERCA uncoupler, THADA is involved in the control of thermogenesis and energy production.

Compared to other eutherian mammals, naked-mole rats show an extraordinary biology e.g. related to unusual thermogenesis and longevity (Ruby et al., 2018; Dammann et al., 2019). The former prompted us to perform an in-depth analysis of NMRs Thada in comparison with other mammals. This comparison led to the identification of four amino acid exchanges in NMR at positions where all other mammals analyzed share the same amino acids. In addition, NMRs Thada shows one deletion not seen in any of the other mammals investigated.

In Drosophila, *THADA* knockout leads to obese hyperphagic flies with reduced energy production and enhanced cold-sensitivity (Muraru et al., 2017). These effects could be reversed by transfer of intact human *THADA*. As a next experimental step an in depth characterization of the NMR-unique changes of THADA as determined herein seems reasonable. It e.g. remains to be determined if transfection of NMR’s *Thada* and *Thada* with single changes as present in NMR can also reverse these effects. In order to analyze the possible biological significance of the NMR-unique mutations NMR-derived fibroblasts can also be transfected using human THADA to study the effects on the transcriptome.

## References

Bai H, Liu H, Suyalatu S, Guo X, Chu S, Chen Y, et al. Association Analysis of Genetic Variants with Type 2 Diabetes in a Mongolian Population in China. J Diabetes Res. 2015;2015:613236. https://www.hindawi.com/journals/jdr/2015/613236/

Bakhashab S, Ahmed N. Genotype based Risk Predictors for Polycystic Ovary Syndrome in Western Saudi Arabia. Bioinformation. 2019 Dec 10;15(11):812–819. http://www.bioinformation.net/015/97320630015812.htm

Bol S, Belge G, Rippe V, Bullerdiek J. Molecular cytogenetic investigations define a subgroup of thyroid adenomas with 2p21 breakpoints clustered to a region of less than 450 kb. Cytogenet Cell Genet. 2001;95(3-4):189–91. doi: 10.1159/000059344. https://www.karger.com/Article/Abstract/59344

Boesgaard TW, Gjesing AP, Grarup N, Rutanen J, Jansson PA, Hribal ML, et al. Variant near ADAMTS9 known to associate with type 2 diabetes is related to insulin resistance in offspring of type 2 diabetes patients--EUGENE2 study. PLoS One. 2009 Sep 30;4(9):e7236. https://journals.plos.org/plosone/article?id=10.1371/journal.pone.0007236

Braude S, Holtze S, Begall S, Brenmoehl J, Burda H, Dammann P, et al. Surprisingly long survival of premature conclusions about naked mole-rat biology. Biol Rev Camb Philos Soc. 2021 Apr;96(2):376–393. https://onlinelibrary.wiley.com/doi/10.1111/brv.12660

Cardona A, Pagani L, Antao T, Lawson DJ, Eichstaedt CA, Yngvadottir B, et al. Genome-wide analysis of cold adaptation in indigenous Siberian populations. PLoS One. 2014 May 21;9(5). https://journals.plos.org/plosone/article?id=10.1371/journal.pone.0098076

Chatterjee N, Perrimon N. Thermogenesis by THADA. Dev Cell. 2017 Apr 10;41(1):1–2. https://www.cell.com/developmental-cell/fulltext/S1534-5807(17)30207-1?_returnURL= https://linkinghub.elsevier.com%2Fretrieve%2Fpii%2FS1534580717302071%3Fshowall%3Dtrue

Cheng L, Caberto CP, Lum-Jones A, Seifried A, Wilkens LR, Schumacher FR, et al. Type 2 diabetes risk variants and colorectal cancer risk: the Multiethnic Cohort and PAGE studies. Gut. 2011 Dec;60(12):1703–11. https://gut.bmj.com/content/60/12/1703

Cui L, Zhao H, Zhang B, Qu Z, Liu J, Liang X, et al. Genotype-phenotype correlations of PCOS susceptibility SNPs identified by GWAS in a large cohort of Han Chinese women. Hum Reprod. 2013 Feb;28(2):538–44. https://academic.oup.com/humrep/article/28/2/538/599478

Cui L, Li G, Zhong W, Bian Y, Su S, Sheng Y, Shi Y, et al. Polycystic ovary syndrome susceptibility single nucleotide polymorphisms in women with a single PCOS clinical feature. Hum Reprod. 2015 Mar;30(3):732–6. https://academic.oup.com/humrep/article/30/3/732/662076

Dammann P, Scherag A, Zak N, Szafranski K, Holtze S, Begall S, et al. Comment on ‘Naked mole-rat mortality rates defy Gompertzian laws by not increasing with age’. Elife. 2019 Jul 9;8:e45415. https://elifesciences.org/articles/45415

Day FR, Hinds DA, Tung JY, Stolk L, Styrkarsdottir U, Saxena R, et al. Causal mechanisms and balancing selection inferred from genetic associations with polycystic ovary syndrome. Nat Commun. 2015 Sep 29;6. https://www.nature.com/articles/ncomms9464

DeMenna J, Puppala S, Chittoor G, Schneider J, Kim JY, Shaibi GQ, et al. Association of common genetic variants with diabetes and metabolic syndrome related traits in the Arizona Insulin Resistance registry: a focus on Mexican American families in the Southwest. Hum Hered. 2014;78(1):47–58. https://www.karger.com/Article/Abstract/363411

Drieschner N, Kerschling S, Soller JT, Rippe V, Belge G, Bullerdiek J, et al. A domain of the thyroid adenoma associated gene (THADA) conserved in vertebrates becomes destroyed by chromosomal rearrangements observed in thyroid adenomas. Gene. 2007 Nov 15;403(1–2):110–7. https://www.sciencedirect.com/science/article/abs/pii/S0378111907003769?via%3Dihub

Eriksen MB, Brusgaard K, Andersen M, Tan Q, Altinok ML, Gaster M, Glintborg D. Association of polycystic ovary syndrome susceptibility single nucleotide polymorphism rs2479106 and PCOS in Caucasian patients with PCOS or hirsutism as referral diagnosis. Eur J Obstet Gynecol Reprod Biol. 2012 Jul;163(1):39–42. https://www.ejog.org/article/S0301-2115(12)00140-6/fulltext

García-Casas P, Pilar Alvarez-Illera, Fonteriz RI, Montero M, Alvarez J. Mechanism of the lifespan extension induced by submaximal SERCA inhibition in C. elegans. Mech Ageing Dev 2021, 196, 111474 https://www.sciencedirect.com/science/article/abs/pii/S0047637421000464?via%3Dihub

Goodarzi MO, Jones MR, Li X, Chua AK, Garcia OA, Chen YD, et al. Replication of association of DENND1A and THADA variants with polycystic ovary syndrome in European cohorts. J Med Genet. 2012 Feb;49(2):90–5. https://jmg.bmj.com/content/49/2/90

Green RE, Krause J, Briggs AW, Maricic T, Stenzel U, Kircher M, et al. A draft sequence of the neandertal genome. Science (80-). 2010 May 7;328(5979):710–22. https://science.sciencemag.org/content/328/5979/710

Grarup N, Andersen G, Krarup NT, Albrechtsen A, Schmitz O, Jørgensen T, et al. Association testing of novel type 2 diabetes risk alleles in the JAZF1, CDC123/CAMK1D, TSPAN8, THADA, ADAMTS9, and NOTCH2 loci with insulin release, insulin sensitivity, and obesity in a population-based sample of 4,516 glucose-tolerant middle-aged Danes. Diabetes. 2008 Sep;57(9):2534–40. https://diabetes.diabetesjournals.org/content/57/9/2534.long

Gupta V, Vinay D G, Rafiq S, Kranthikumar M V, Janipalli C S, Giambartolomei C, et al. Association analysis of 31 common polymorphisms with type 2 diabetes and its related traits in Indian sib pairs. Diabetologia. 2012 Feb;55(2):349–57. https://link.springer.com/article/10.1007%2Fs00125-011-2355-6

Ha L, Shi Y, Zhao J, Li T, Chen ZJ. Association Study between Polycystic Ovarian Syndrome and the Susceptibility Genes Polymorphisms in Hui Chinese Women. PLoS One. 2015 May 15;10(5):e0126505. https://journals.plos.org/plosone/article?id=10.1371/journal.pone.0126505

Hotta K, Kitamoto A, Kitamoto T, Mizusawa S, Teranishi H, So R, et al. Association between type 2 diabetes genetic susceptibility loci and visceral and subcutaneous fat area as determined by computed tomography. J Hum Genet. 2012 May;57(5):305–10. https://www.nature.com/articles/jhg201221

Hu C, Zhang R, Wang C, Wang J, Ma X, Lu J, et al. PPARG, KCNJ11, CDKAL1, CDKN2A-CDKN2B, IDE-KIF11-HHEX, IGF2BP2 and SLC30A8 are associated with type 2 diabetes in a Chinese population. PLoS One. 2009 Oct 28;(10):e7643. https://journals.plos.org/plosone/article?id=10.1371/journal.pone.0007643

Kang ES, Kim MS, Kim CH, Nam CM, Han SJ, Hur KY, et al. Association of common type 2 diabetes risk gene variants and posttransplantation diabetes mellitus in renal allograft recipients in Korea. Transplantation. 2009 Sep 15;88(5):693–8. https://journals.lww.com/transplantjournal/Fulltext/2009/09150/Association_of_Common_Type_2_Diabetes_Risk_Gene.14.aspx

Kloth L, Belge G, Burchardt K, Loeschke S, Wosniok W, Fu X, Nimzyk R, Mohamed SA, Drieschner N, Rippe V, Bullerdiek J. Decrease in thyroid adenoma associated (THADA) expression is a marker of dedifferentiation of thyroid tissue. BMC Clin Pathol. 2011 Nov 4;11:13. https://bmcclinpathol.biomedcentral.com/articles/10.1186/1472-6890-11-13

Morariu EM, McCoy KL, Chiosea SI, Nikitski AV, Manroa P, Nikiforova MN, Nikiforov YE. Clinicopathologic Characteristics of Thyroid Nodules Positive for the THADA-IGF2BP3 Fusion on Preoperative Molecular Analysis. Thyroid. 2021 Aug;31(8):1212–1218. https://www.liebertpub.com/doi/10.1089/thy.2020.0589

Moraru A, Cakan-Akdogan G, Strassburger K, Males M, Mueller S, Jabs M, et al. THADA Regulates the Organismal Balance between Energy Storage and Heat Production. Dev Cell. 2017 Apr 10;41(1):72-81.e6. https://www.cell.com/developmental-cell/fulltext/S1534-5807(17)30169-7?_returnURL= https://linkinghub.elsevier.com%2Fretrieve%2Fpii%2FS1534580717301697%3Fshowall%3Dtrue

Panebianco F, Kelly LM, Liu P, Zhong S, Dacic S, Wang X, et al. THADA fusion is a mechanism of IGF2BP3 activation and IGF1R signaling in thyroid cancer. Proc Natl Acad Sci U S A. 2017 Feb 28;114(9):2307–2312. https://www.pnas.org/content/114/9/2307

Park S, Liu M, Zhang T. THADA_rs13429458 Minor Allele Increases the Risk of Polycystic Ovary Syndrome in Asian, but Not in Caucasian Women: A Systematic Review and Meta-Analysis. Horm Metab Res. 2019 Oct;51(10):661–670. https://www.thieme-connect.de/products/ejournals/abstract/10.1055/a-0969-1872

Pool C, Walter V, Bann D, Goldenberg D, Broach J, Hennessy M, et al. Molecular characterization of tumors meeting diagnostic criteria for the non-invasive follicular thyroid neoplasm with papillary-like nuclear features (NIFTP). Virchows Arch. 2019 Mar;474(3):341–351. https://link.springer.com/article/10.1007%2Fs00428-018-02512-6

Prasad RB, Lessmark A, Almgren P, Kovacs G, Hansson O, Oskolkov N, et al. Excess maternal transmission of variants in the THADA gene to offspring with type 2 diabetes. Diabetologia. 2016 Aug;59(8):1702–13. https://link.springer.com/article/10.1007/s00125-016-3973-9

Rippe V, Drieschner N, Meiboom M, Escobar HM, Bonk U, Belge G, et al. Identification of a gene rearranged by 2p21 aberrations in thyroid adenomas. Oncogene. 2003 Sep 4;22(38):6111–4.

Ruby JG, Smith M, Buffenstein R. Naked Mole-Rat mortality rates defy gompertzian laws by not increasing with age. Elife. 2018 Jan 24;7:e31157. https://elifesciences.org/articles/31157

Ruby JG, Smith M, Buffenstein R. Response to comment on ‘Naked mole-rat mortality rates defy Gompertzian laws by not increasing with age’. Elife. 2019 Jul 9;8:e47047. https://elifesciences.org/articles/47047

Staiger H, Machicao F, Kantartzis K, Schäfer SA, Kirchhoff K, Guthoff M, et al. Novel meta-analysis-derived type 2 diabetes risk loci do not determine prediabetic phenotypes. PLoS One. 2008 Aug 20;3(8):e3019. https://journals.plos.org/plosone/article?id=10.1371/journal.pone.0003019

Schleinitz D, Tönjes A, Böttcher Y, Dietrich K, Enigk B, Koriath M, et al. Lack of significant effects of the type 2 diabetes susceptibility loci JAZF1, CDC123/CAMK1D, NOTCH2, ADAMTS9, THADA, and TSPAN8/LGR5 on diabetes and quantitative metabolic traits. Horm Metab Res. 2010 Jan;42(1):14–22. https://www.thieme-connect.de/products/ejournals/abstract/10.1055/s-0029-1233480

Simonis-Bik AM, Nijpels G, Van Haeften TW, Houwing-Duistermaat JJ, Boomsma DI, Reiling E, et al. Gene variants in the novel type 2 diabetes loci CDC123/CAMK1D, THADA, ADAMTS9, BCL11A, and MTNR1B affect different aspects of pancreatic β-cell function. Diabetes. 2010 Jan;59(1): 293–301. https://diabetes.diabetesjournals.org/content/59/1/293

Soller JT, Beuing C, Escobar H, Winkler S, Reimann-Berg N, Drieschner N, et al. Chromosomal assignment of canine THADA gene to CFA 10q25. Mol Cytogenet [Internet]. 2008 [cited 2021 Jan 28];1(1):11. https://molecularcytogenetics.biomedcentral.com/articles/10.1186/1755-8166-1-11

Sun M, Sheng Y, Ma Z, Chen Z, Tang R. [Correlation analysis between polycystic ovary syndrome susceptibility genes and metabolic phenotypes]. Zhonghua Fu Chan Ke Za Zhi. 2014 Jun;49(6):441–5.

Tian Y, Li J, Su S, Cao Y, Wang Z, Zhao S, Zhao H. PCOS-GWAS Susceptibility Variants in THADA, INSR, TOX3, and DENND1A Are Associated With Metabolic Syndrome or Insulin Resistance in Women With PCOS. Front Endocrinol (Lausanne). 2020 Apr 30;11:274. https://www.frontiersin.org/articles/10.3389/fendo.2020.00274/full

Villegas R, Goodloe RJ, McClellan BE Jr, Boston J, Crawford DC. Gene-carbohydrate and gene-fiber interactions and type 2 diabetes in diverse populations from the National Health and Nutrition Examination Surveys (NHANES) as part of the Epidemiologic Architecture for Genes Linked to Environment (EAGLE) study. BMC Genet. 2014 Jun 14;15:69. https://bmcgenomdata.biomedcentral.com/articles/10.1186/1471-2156-15-69

Vishnubotla DS, Shek AP, Madireddi S. Pooled genetic analysis identifies variants that confer enhanced susceptibility to PCOS in Indian ethnicity. Gene. 2020 Aug 20;752:144760. https://www.sciencedirect.com/science/article/abs/pii/S0378111920304297?via%3Dihub

Xia JY, Tian W, Yin GH, Yan H. Association of Rs13405728, Rs12478601, and Rs2479106 single nucleotide polymorphisms and in vitro fertilization and embryo transfer efficacy in patients with polycystic ovarian syndrome: A case control genome-wide association study. Kaohsiung J Med Sci. 2019 Jan;35(1):49–55. https://onlinelibrary.wiley.com/doi/10.1002/kjm2.12008

West HL, Corbin KL, D’Angelo CV, Donovan LM, Jahan I, Gu G, Nunemaker CS. Postnatal maturation of calcium signaling in islets of Langerhans from neonatal mice. Cell Calcium. 2021 Mar;94:102339. https://www.sciencedirect.com/science/article/abs/pii/S0143416020301810?via%3Dihub

Zhao H, Xu X, Xing X, Wang J, He L, Shi Y, et al. Family-based analysis of susceptibility loci for polycystic ovary syndrome on chromosome 2p16.3, 2p21 and 9q33.3. Hum Reprod. 2012 Jan;27(1):294–8. https://academic.oup.com/humrep/article/27/1/294/717131

